# Wild juvenile Arctic char (*Salvelinus alpinus*) show seasonal acclimatization of cardiac and whole-animal heat tolerance

**DOI:** 10.64898/2026.11.10.764389

**Authors:** Shira A. Berkelhammer, Les N. Harris, Emily P. Williams, Gabriel Ferland, Bonnie M. Hamilton, Jean-Sébastien Moore, Matthew J.H. Gilbert

## Abstract

Arctic fishes experience extreme seasonality, yet how seasonal acclimatization shapes their ability to cope with thermal variation remains poorly understood. The Arctic char (*Salvelinus alpinus*) is one Arctic fish of immense cultural, subsistence, and economic value and encounters temperatures ranging from below 0°C to above 21°C. Thermal physiology commonly varies across life stages, but has been largely unexamined in wild juvenile Arctic char. We performed winter and summer field-based assessments of wild juvenile Arctic char thermal habitat use, whole-animal and cardiac heat tolerance, and mechanisms shaping the acute thermal sensitivity of heart rate (*f*_H_). Average body temperature increased from 1.2°C in winter to 8.9°C in summer, and critical thermal maximum (CT_max_) increased correspondingly from 21.4 to 25.9°C. We assessed the intrinsic and maximum *f*_H_ (*f*_Hint_ and *f*_Hmax_) of anaesthetized fish during acute warming using cholinergic blockade (atropine) and either β-adrenergic blockade (sotalol; *f*_Hint_) or stimulation (isoproterenol; *f*_Hmax_). *f*_H_ increased with acute warming in both seasons and treatments. However, in summer, *f*_H_ was more responsive to warming (Q_10_ from 5 to 15°C: 2.4 vs. 2.0 in winter), reached a higher peak, and continued to increase while maintaining rhythmicity up to higher temperatures, with the onset of heat-induced cardiac arrhythmia (T_arr_) occurring at 23.6°C vs. 18.7°C in winter. Contrary to expectations, β-adrenergic blockade had little effect on these cardiac parameters. During acute warming, winter-acclimatized fish exhibited slower atrioventricular conduction (longer PR interval) and slower ventricular depolarization (longer QRS duration), which may underlie their lower cardiac heat tolerance and the reduced responsiveness of *f*_H_ to warming. Combined, our results show that wild Arctic char exhibit strong seasonal plasticity of acute thermal performance, demonstrating the value of field-based physiological assessments and the need to consider plasticity in evaluating responses to environmental change.

## Introduction

Arctic fishes commonly contend with extreme thermal variation, spending most of their lives in near-freezing water beneath the ice but facing summer temperatures that can exceed 20°C in freshwater systems (Gilbert *et al*., 2016, 2020; Dubos *et al*., 2025). Beyond these substantial seasonal changes, further thermal variation is encountered as fish migrate throughout their environment and in response to stochastic weather events. Fish in Arctic lakes often lack opportunities to reduce this variation through behavioral thermoregulation because many Arctic lakes do not thermally stratify, thus limiting access to thermal refugia during heat events (Dubos *et al*., 2025). Additionally, seasonal light varies drastically in the far north with no daylight in the depths of winter and for 24 hours for parts of the summer. Replicating such environmental variation in a laboratory setting is challenging and few studies have characterized the physiological plasticity associated with such extreme seasonality in Arctic fishes (Gilbert and Farrell, 2021; Middleton *et al*., 2024; Procopio *et al*., 2024; Gilbert *et al*., 2026), including wild individuals. Given the logistical, financial and physical challenges of winter and northern research, there is a particular lack of information on winter physiology of northern fishes (Farrell *et al*., 2003; Gilbert and Tierney, 2018; Mallory *et al*., 2018).

The heart is one integral organ that commonly exhibits pronounced thermal and seasonal plasticity, including in Arctic fishes (Abramochkin and Vornanen, 2015; Keen *et al*., 2017; Gilbert and Farrell, 2021). A progressive increase in temperature drives an increase in tissue oxygen demand, requiring a greater cardiac output. Stroke volume commonly does not increase with warming until extreme temperatures (Gollock *et al*., 2006; Farrell *et al*., 2009; Ekström *et al*., 2014; Penney *et al*., 2014; Gamperl *et al*., 2022; but see El *et al*., 2025), and so this increase in cardiac output is met, in large part, by increasing heart rate (*f*_H_); *f*_H_ increases up to a warm-induced peak (peak*f*_H_) and then typically plateaus or declines with further warming until the heartbeat loses rhythmicity causing a collapse in *f*_H_ (Figure S1, Anttila *et al*., 2014; Vornanen, 2016; Haverinen and Vornanen, 2020; Gilbert *et al*., 2024; Porter and Gamperl, 2026). Given the centrality of *f*_H_ in meeting temperature dependent oxygen demands, it is not surprising that fishes commonly exhibit plasticity in *f*_H_ and its acute response to thermal variation as part of thermal acclimation (Anttila *et al*., 2014, 2015; Gilbert and Farrell, 2021; Gilbert *et al*., 2022b, 2022a; Strowbridge *et al*., 2025). Cardiac thermal plasticity includes modification of cardiac pacemaker rates, the electrical excitability of the heart, and the role of adrenaline in controlling those processes (Haverinen and Vornanen, 2004, 2007, 2009; Abramochkin and Vornanen, 2015; Vornanen, 2016; Sutcliffe *et al*., 2020; Williams *et al*., 2025). However, seasonal acclimatization includes complex responses to a broad range of environmental conditions and seasonal endogenous cycles. Thus, seasonal responses in the wild do not always align with what would be expected from thermal acclimation alone (Filatova *et al*., 2019). Our understanding of these relationships is particularly lacking in Arctic fishes, and in general of wild fish in their natural habitat.

The Arctic char is one iconic Arctic fish species that displays remarkable cardiac thermal acclimation potential (Gilbert *et al*., 2020, 2022b; Gilbert and Farrell, 2021), however cardiac seasonal plasticity has yet to be examined. Presently, the thermal physiology of Arctic char has been studied in wild anadromous adults (Gilbert *et al*., 2020, 2022b), hatchery/laboratory raised juveniles (∼1.3-18cm: Baroudy and Elliott, 1994; ∼25g: Anttila *et al*., 2015) and intermediately (∼400 g) sized individuals (Gilbert and Farrell, 2021). For example, in Gilbert and Farrell (2021), lab-raised Arctic char acclimated to 14°C (a warm summer temperature) reset their maximum *f*_H_ (*f*_Hmax_) and showed strong compensation of heart rate and cardiac heat tolerance that was 35-45% higher than the 2°C winter acclimated fish. In a parallel summer field study on wild anadromous adult migrating Arctic char, Gilbert *et al*. (2020) showed that *f*_Hmax_ and the temperature that induces peak *f*_H_ (T_peak_) were comparable to Arctic char raised at 6 and 10°C in the lab. However, data are not available for winter acclimatized or juvenile Arctic char. Thermal physiology commonly varies with season, fish body mass, and life stage, with juveniles often having greater acute heat tolerance and higher metabolic rates than mature adults relative to their body mass (Filatova *et al*., 2019; Mayer *et al*., 2024). Moreover, juveniles remain in freshwater habitats until they undergo spring migration at about 3-6 years old, further differentiating themselves from adults (Gilbert *et al*., 2016). There is reason to expect that seasonal plasticity may not simply reflect thermal acclimation because, in addition to temperature, Arctic char face strong seasonal variation in dissolved oxygen, food availability, photoperiod, and annual endogenous cycles (Jørgensen *et al*., 2013; Hawley *et al*., 2017; Dubos *et al*., 2025, 2026). In addition to changes in *f*_H_ and cardiac heat tolerance, the adrenergic control of cardiac function and its sensitivity to temperature can vary seasonally and with thermal acclimation, but this plasticity varies greatly among species and has not been examined in Arctic char (Keen *et al*., 1993; Aho and Vornanen, 2001; Shiels *et al*., 2003; Eliason *et al*., 2011; Williams *et al*., 2025).

Given the knowledge gaps highlighted above, our aim was to determine 1) how cardiac thermal sensitivity and heat tolerance change seasonally, 2) to what extent these changes are driven by differences in adrenergic scope for heart rate, and 3) how these changes relate to changes in whole-animal acute heat tolerance in wild, juvenile Arctic char. We hypothesized that seasonally-acclimatized juvenile Arctic char would show classic thermal compensation of heart rate and heat tolerance. Accordingly, we predicted that winter-acclimatized juvenile Arctic char would show relatively high heart rates at low to moderate temperatures to counteract the depressive effect of cold, but that they would be less able to maintain or elevate heart rate at higher temperatures compared to summer-acclimatized fish.

## Methods

Sampling occurred at Inuhuktok (69°14’16”N, 104°45’41”W), an Arctic char spawning lake near Ikaluktutiak, Cambridge Bay NU, Canada (Figure 1, Table 1). Field work took place during May and August 2024-2025, in which May was classified as “winter” and August as “summer”. Juvenile Arctic char (Table 1) were caught at the end of winter (May) and summer (August) with 50 m 1 ½ and 2-inch (stretched measure) gillnets. Fish were confirmed to be juveniles as they fell in the size range typical of smolts seen in the region (Gilbert *et al*., 2016). In winter, nets were set through ∼1.9-2.1m of ice using a remotely operated vehicle (ROV: Chasing M2; Chasing Innovation Ltd), a method that took two winter field seasons to develop. The ROV was deployed under the ice through a larger net hole (∼100 x 70cm) to retrieve a pull line from a smaller (25 cm diameter) rope hole ∼50 m away. The pull line was then used to deploy the net. In summer, nets were anchored to shore and set by boat. Following capture, fish were transferred to a mesh holding pen (∼1 m^3^) in the summer (∼7.4°C) or to a temperature-controlled, aerated cooler during winter (∼0.7°C) and held for up to 24 h with dissolved oxygen monitoring before tagging or experimentation. Sampling and animal use protocols were approved by the University of Alaska Fairbanks Institutional Animal Care and Use Committee ( # 2181369-3, # 2302455-1) and Fisheries and Oceans Canada (OPA ACC-2024-40, OPA-ACC-2025-3).

**Figure 1.**
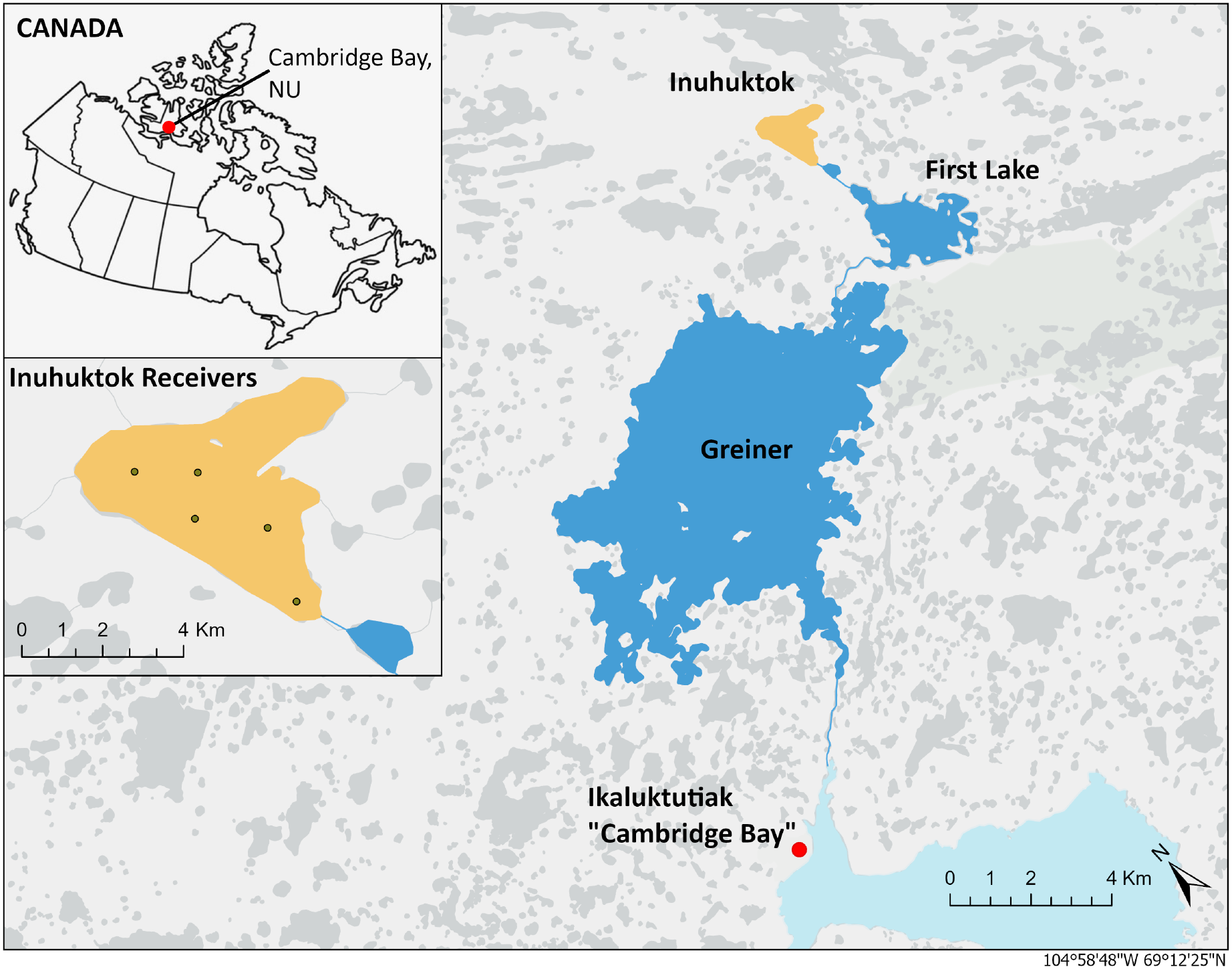
Location of the study area near Cambridge Bay (red dot), in the Kitikmeot Region of Nunavut, Canada. Inuhuktok, the study lake, is highlighted in yellow, and the locations of five acoustic telemetry receivers are shown in the enlarged inset.

**Table 1.** Total sample sizes (n), body mass, fork length, Fulton’s condition factor, and relative ventricular mass of juvenile Arctic char acclimatized to winter (2024 and 2025) and summer (2024) seasonal conditions. Data presented as average ± SEM. Summer 2024 had 3 fish removed from both **f**_Hmax_ and **f**_Hint_ treatment groups, one fish removed from winter 2024 **f**_Hmax_, and one fish removed from winter 2025 CT_max_.

| **Season** | **Treatment** | ***n*** | **Body mass (g)** | **Length (mm)** | **Fulton’s condition factor** | **Relative ventricular mass (% body mass)** |
|---|---|---|---|---|---|---|
| Winter | *f*<sub>Hmax</sub> | 12 | 70.1 ± 5.0 | 198.3 ± 5.6 | 0.89 ± 0.03 | 0.12 ± 0.01 |
|  | *f*<sub>Hint</sub> | 9 | 68.8 ± 7.3 | 199.6 ± 7.5 | 0.85 ± 0.04 | 0.10 ± 0.01 |
|  | CT<sub>max</sub> | 11 | 57.0 ± 2.5 | 193.0 ± 2.9 | 0.79 ± 0.01 | – |
|  | Tagging | 10 | 79.9 ± 4.1 | 209.6 ± 3.7 | 0.90 ± 0.02 | – |
| Summer | *f*<sub>Hmax</sub> | 9 | 137.2 ± 12.9 | 234.8 ± 6.2 | 1.03 ± 0.04 | 0.13 ± 0.01 |
|  | *f*<sub>Hint</sub> | 9 | 149.5 ± 10.2 | 241.9 ± 5.5 | 1.04 ± 0.03 | 0.13 ± 0.01 |
|  | CT<sub>max</sub> | 12 | 58.9 ± 2.9 | 190.3 ± 3.5 | 0.90 ± 0.02 | – |
|  | Tagging | 15 | 236.5 ± 41.3 | 276.5 ± 13.8 | 1.0 ± 0.03 | – |

### Environmental and body temperature

We measured water and fish body temperature using environmental loggers and surgically implanted acoustic transmitters, respectively. Following the protocol of Dubos et al. (2025), five acoustic receivers (Innovasea VR2AR, 69 kHz) were deployed in Inuhuktok from August 2023 to August 2025 (Figure 1). One receiver mooring was equipped with two dissolved oxygen loggers (miniDOT, PME, USA, 18.7 m and 2 m from bottom anchor) and two temperature loggers (HOBO 64K Pendant, Onset, USA, 18.7 m and 12.5 m above the anchor) to characterize environmental conditions at different depths.

Juvenile Arctic char were surgically implanted with V7 temperature-pressure acoustic transmitters (tags) (V7TP-2x, diameter: 7mm, length: 22 mm, weight: 1.5 g, Innovasea Ltd.) in August 2024 and May 2025. A longer nominal transmission delay of 300s was selected to extend transmitter battery life. Fish suitability for tagging was assessed based on the severity of gillnet-related injuries and overall activity levels. Suitable fish were then anesthetized using tricaine methanesulfonate (150 mg L ^-1^ TMS; Syndel, Nanaimo, Canada) to level III anesthesia (CCAC: Canadian Council on Animal Care guidelines), measured for weight and fork length, and placed in a V-shaped surgical cradle. During surgery, gills were continuously irrigated with water containing 75 mg L^-1^ TMS to maintain anesthesia. A small incision was made on the ventral surface of the fish, posterior to the pelvic fins to insert the transmitter into the body cavity. The incision was closed using 5-0 coated Vicryl sutures (Ethicon, J463G). A pectoral fin clip was collected and preserved in ethanol for subsequent genetic sex determination and fish were transferred to a recovery cooler containing fresh lake water. Once fully recovered, fish were released back into the lake.

Each receiver in the lake recorded ambient water temperature, in addition to the external temperature probes deployed in the lake. Body temperature data were obtained from acoustic detections of the implanted temperature-sensing transmitters recorded by the receivers. In total, the dataset comprised two years of environmental data spanning August 2023 to August 2025, with one year of body temperature and telemetry data. Fish were implanted with acoustic transmitters in August 2024 and the receivers retrieved August 2025. Body temperature and telemetry data were recorded throughout this period, except between January and May 2025, when transmitter batteries were depleted, resulting in a gap in detections.

### Critical thermal maximum

Whole-animal thermal tolerance was assessed as the critical thermal maximum (CT_max_). Measuring CT_max_ is a useful indicator of upper bounds for thermal performance, and thus whole-organism heat tolerance, and can be valuable for comparing between and within fish species with different rearing histories (Sandblom *et al*., 2016). In 2025, juveniles were transferred to a well-aerated 100 L holding cooler containing one submersible 800W titanium aquarium heater (Finnex, Countryside, IL) and 30 L of water. Fish were held at ambient water temperature for 30 min, and then acutely warmed at 6°C h^-1^ up to the temperature at which they could no longer maintain dorsoventral equilibrium (Beitinger *et al*., 2000; Morgan *et al*., 2018; Gilbert *et al*., 2020). Test temperature for winter acclimatized fish (n = 11) began at 2°C, and for summer acclimatized fish (n =12) at 10°C. One winter individual was excluded from analysis due to a netting injury during capture. Following completion of the trial, fish were euthanized, weight and length were measured, and tissue samples were taken for related projects.

### Heart rate assessment

Cardiac thermal tolerance was evaluated in both seasons in 2024 and winter 2025 by measuring heart rate in anaesthetized juveniles fitted with sub-dermal electrocardiogram (ECG) electrodes. This method has been previously developed for and validated in salmonids, including Arctic char (Casselman *et al*., 2012; Anttila *et al*., 2014; Gilbert and Farrell, 2021; Gilbert *et al*., 2024) and has its inherent strengths and limitations which have been previously discussed at length (Gilbert *et al*., 2024). Individual fish were anesthetized in 150 mg L ^-1^ buffered TMS (1:2 sodium bicarbonate) in freshwater at 2°C (winter) and 10°C (summer) until ventilation slowed but did not stop (Anesthesia Stage III CCAC; Gilbert *et al*., 2024). Fish were weighed, measured for fork length, and transferred to mesh cradle in an anesthetic bath containing a lower maintenance dosage of 70 mg L ^-1^ buffered TMS. Fish were placed supine and fitted with individual pumps to continuously irrigate their gills (Gilbert *et al*., 2024; Williams *et al*., 2025). Four fish were evaluated simultaneously, and each individual fish was considered an independent experimental unit. The anesthetic bath was set at 2°C for winter acclimatized fish and 5°C for summer acclimatized fish, using a flow through aquarium chiller (TK 2000, Teco, Ravenna, ITA). Summer experiments were started at 5°C to avoid cold-induced arrhythmia that was witnessed in preliminary summer 2024 trials as we were optimizing our methods. Stainless steel electrodes were inserted sub-dermally on the ventral surface directly over the heart and just above the pelvic fins, with a reference electrode placed in the mesh cradle. The ECG signal was collected through BioAmp amplifiers (one 4-channel DP-304A and two single-channel DP-301 differential amplifiers; ADInstruments, Sydney, Australia) connected to a PowerLab 8/35 data acquisition unit. Signals were amplified and filtered before being recorded in LabChart software (version 8.1.25). Water temperature near the gill outflow was continuously monitored using a T-type thermocouple pod (ADInstruments) linked to the PowerLab, along with a handheld conductivity, temperature and oxygen sensor (ODO Conductivity/Temperature probe, ProSolo, Xylem Inc., Ohio, USA). The temperature probe linked to the PowerLab experienced periodic calibration issues, so the handheld probe was used to continuously verify temperature. The final temperature values used came from the conductivity, temperature, and oxygen sensor.

To assess the role of adrenaline, each fish received an intraperitoneal injection of atropine sulphate (1.2 mg kg^-1^; Sigma-Aldrich, St. Louis, USA) and an injection of either sotalol (2.7 mg kg^-1^; Sigma-Aldrich) or isoproterenol (4 μg kg^-1^; Sigma-Aldrich) saline solutions (0.8% NaCl) in a total volume of 1 mL kg^-1^ to determine *f*_Hint_ (winter: n=9, summer: n=12) and *f*_Hmax_ (winter: n = 13, summer: n=12), respectively. Atropine sulphate acts as a muscarinic acetylcholine receptor antagonist, blocking parasympathetic inhibition of *f*_H_, while isoproterenol functions as a β-adrenoreceptor agonist and sotalol as a β-adrenergic antagonist. Once fish received the injections, 20 minutes were given for the drugs to take effect and for heart rate to stabilize before beginning the experiment. Temperature was then raised at a rate of 6°C h^-1^ in 1°C increments using a submersible 800W titanium aquarium heater until cardiac arrhythmia occurred for each fish. After all fish had gone arrhythmic, they were sampled for body mass, sex, fat %, carcass weight, gonad weight, ventricle weight, gut weight, pylorus weight, liver weight, and intestine weight and length. In total, seven fish were removed from the final *f*_H_ dataset. This included one fish from winter 2024, and 6 fish from summer 2024, with 4 of the 6 removed due to cold shock arrhythmia. All other fish were removed due to electrical interference that disrupted data collection leaving winter **f**_Hmax_ n=12, **f**_Hint_ n=9, and summer **f**_Hmax_ n=9, **f**_Hint_ n=9.

### Data Analysis

#### Electrocardiogram waveform analysis

Heart rate (*f*H) was quantified from ECG recordings using the cyclic measurements feature in the Labchart software. A 30-second segment from the final minute of each 1°C warming step was analyzed with automated beat detection, and the results were visually checked to confirm accuracy. When poor signal quality or electrical noise prevented reliable detection across the full 30 s, the average of 10 manually verified beats was used instead. Temperature values were adjusted based on recorded values to adjust for computer temperature probe calibration issues using a handheld conductivity, temperature and oxygen sensor (Xylem Inc., Ohio, USA). The thermal tolerance metrics determined from the ECG analysis included the warming response of *f*_Hint_ and *f*_Hmax_, peak *f*_H_ reached (peak*f*_H_), temperature at peak *f*_H_ (T_peak_), and temperature at the onset of arrhythmia (T_arr_) were identified for summer and winter acclimatized individuals. Additionally, Δ*f*_H_ was calculated as peak*f*_H_ – *f*_H_ at 5°C for each individual.

Detailed ECG waveform analysis was performed to gain further mechanistic understanding of the seasonal effects of warming on the hearts of juvenile Arctic char. We examined beat intervals from the same segment where *f*_H_ measurements were taken, arranged over 10 beats. To ensure correct and uniform placement across all temperature steps, we used the automatic detection function in the software to identify P, Q, R, S, and T waves and manually adjusted them when necessary. To account for variation seen between the P and T waves, all lines accounting for the P wave (start, peak, end) and the T wave (peak, end) were placed on the respective beat’s peak. The beat intervals PR (the atria-to-ventricle conduction time), RR (the time between two consecutive heartbeats), QT (total duration of electrical activity in the ventricle), and QRS duration (conduction of the electrical impulse across the ventricle) were assessed.

#### Arrhythmia classification

Cardiac arrhythmias were identified as missing or delayed QRS complexes and were initially categorized visually as asystole or atrioventricular (AV) block at the first instance of arrhythmia (Badr *et al*., 2016). Asystole was classified as having a full QRS complex, followed by a clear P-wave and then having no apparent P-waves or QRS complexes for at least one cardiac cycle. Additionally, following this arrhythmia, PR interval to RR interval ratio (PR:RR) did not increase. AV block arrhythmia was initially categorized as type 2 second-degree AV block by site, in which multiple P-waves occurred between one QRS complex, but in which PR:RR ratio remains constant. A type 1 second-degree AV block was characterized by an increase in the PR:RR ratio falling above the 95% confidence interval calculated for that individual’s PR:RR ratio (Vogler *et al*., 2012).

#### Q_10_ value calculations

Incremental, acute, and acclimated Q_10_ values were calculated for *f*_Hmax_ and *f*_Hint_ to assess thermal sensitivity (Anttila *et al*., 2013; Williams *et al*., 2025). Individual fish incremental Q_10_ values were calculated over 2°C temperature change. Acute Q_10_ values were calculated over the acclimatized temperatures of each season (the acute change over 2-10°C in winter and summer fish). Because summer fish *f*_H_ measurements started at 5°C and winter fish at 2°C, acute Q_10_ was calculated between 2-10°C for winter fish only, and over 5-10°C for both summer and winter fish for comparison. Additionally, acute Q_10_ was calculated over 5-15°C as this interval allows us to calculate a Q_10_ over a 10°C range. Acclimated Q_10_ values were calculated between the 2 °C *f*_H_ measurement for winter fish and the 10°C *f*_H_ measurement for summer fish, and 5°C for winter fish and 15°C for summer fish. Data collection spanned two winter field periods (2024 and 2025), but winter 2024 only included *f*_Hmax_ measurements. Therefore, acclimated Q_10_ was calculated for *f*_Hmax_ and *f*_Hint_ between summer 2024 and winter 2025 fish. Acclimated Q_10_ for *f*_Hmax_ was calculated for the following comparisons: winter 2025 vs. summer 2024, winter 2024 vs. summer 2024, and a combined dataset of winter 2025 and summer 2024.

#### Adrenergic scope for heart rate

Adrenergic scope for heart rate (β-adr**f**_H_ scope) was calculated at each temperature step as in Williams et al. 2025, using the equation:

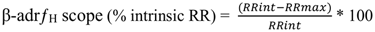

This method considers implications of *f*_H_ overestimating adrenergic tone (Axelsson *et al*., 1992, 2000; Altimiras *et al*., 1997; McKenzie *et al*., 2007; Iversen *et al*., 2010). Tone was calculated by subtracting RRmax (the intervals averaged within a season for maximum *f*_H_ treatments for each temperature) from RRint (intervals averaged within a season for intrinsic *f*_H_ treatments for each temperature) and dividing by RRint. This represents the ability to increase *f*_H_ above the intrinsic rate.

### Statistical Analysis

To examine the effect of seasonal acclimatization, generalized linear mixed effects models (GLMM) were used to analyze *f*_H_, RR interval, PR interval, QT interval, and incremental Q_10_, as they did not meet the assumptions for a linear mixed model (package “glmmTMB”, family=Gamma, link=Log, Brooks *et al*., 2017). The Gamma distribution was used as our data were continuous, positive, and right-skewed. Normality of residuals was assessed using the Shapiro-Wilk test, and homogeneity of variance was evaluated by visual inspection of residuals. By comparison, QRS data met these assumptions and were analyzed using a linear mixed model. All models contained fish ID as a random effect to account for repeated measures, with the parameter of interest modeled as a function of acute temperature, season, and drug treatment (*f*_Hint_ or *f*_Hmax_). Models were selected using the lowest Akaike information criterion (AIC) value. Post-hoc analysis was completed using the R “emmeans” package (Lenth and Piaskowski, 2017).

Seasonal acclimation on upper thermal tolerance metrics was calculated using linear models (LM) as residuals and data of peak*f*_H_, T_peak_, T_arr_, and TQ_10_ was normally distributed. Acute Q_10_ had residuals that were not normally distributed and were assessed using a generalized linear model (GLM; family=Gamma, link=Log). All response variables were modeled as a function of season and drug, with post hoc comparisons done using “emmeans” package with a Holm adjustment. Body mass and relative ventricular mass were only added to models as predictor variables when they lowered AIC values (improving model fit). CT_max_ comparison between winter and summer was calculated using a LM and modeled as a function of season and body mass. Statistical significance was decided at P<0.05, and all values are presented as mean ± standard error (S.E.M) unless otherwise specified. Statistical analyses were completed using R Studio in R (R Core Team, 2014) or Microsoft Excel (v16.99.2, Microsoft Corp., 2024). Figures were prepared using Prism v.10.4.2 (GraphPad Software, San Diego, CA, USA).

## Results

### Environmental and body temperature

Fish body temperature was closely associated with seasonal variation in lake temperature. Fish body temperatures were ∼0.5°C warmer than lake temperatures on average during winter (1.2° vs 0.7 °C, respectively) and 1.5°C warmer during summer (8.9° vs 7.4°C, respectively; Figures 2a, b). The average body temperature during the experimental period in winter 2025 was 1.1°C and 11.1°C during summer 2024 which were used as the estimated acclimatization temperatures throughout.

**Figure 2.**
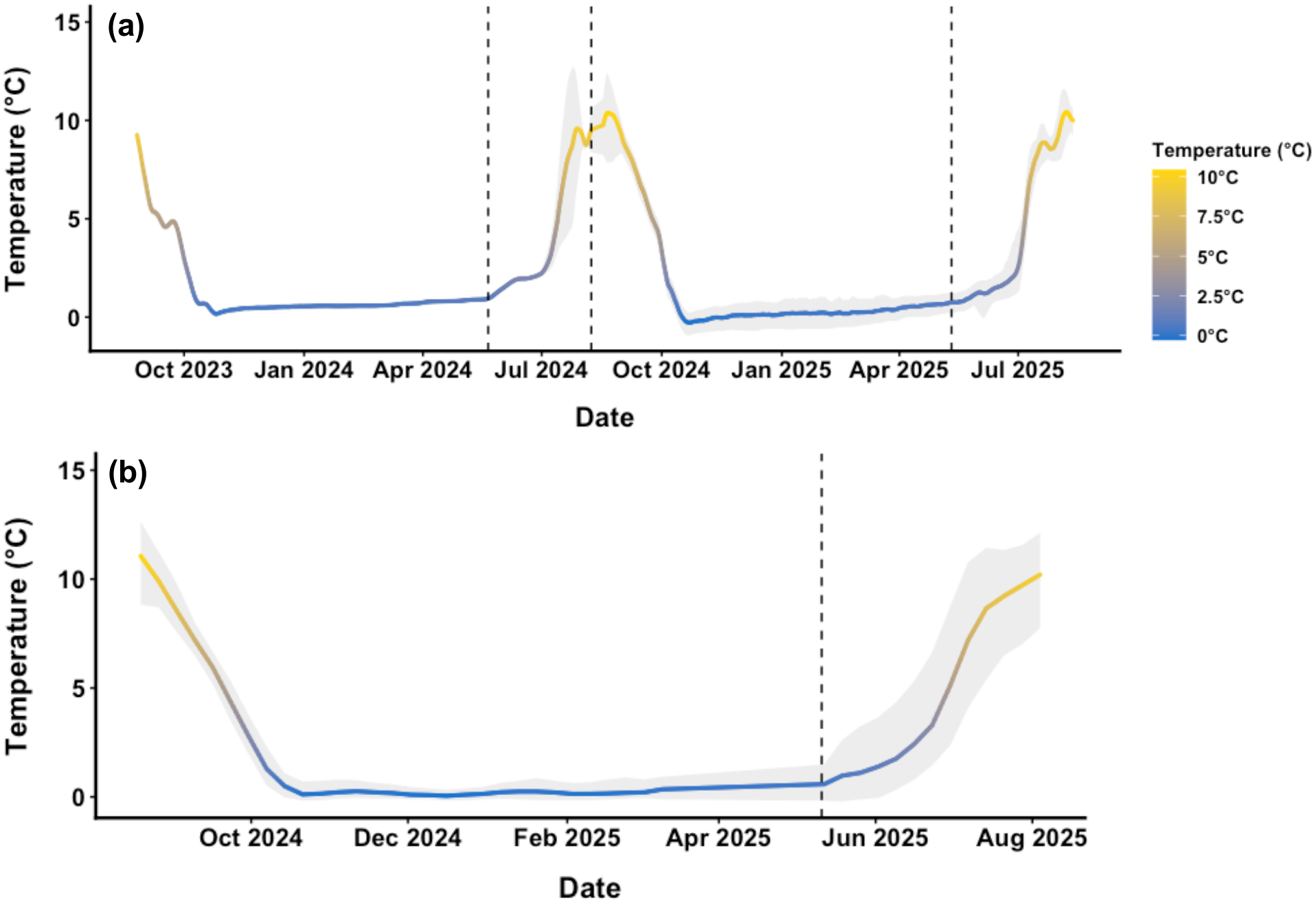
Seasonal variation in (a) water temperature in Inuhuktok from September 2023 to August 2025 and (b) body temperature of juvenile Arctic char from August 2024 through August 2025. Body temperatures were recorded using temperature-sensing acoustic transmitters implanted in fish, whereas lake temperatures were recorded using acoustic receivers. The line in (a) shows the 7-day rolling average of daily temperature averaged across all receiver temperature loggers, shaded area shows the 7-day rolling average of the daily minimum and maximum. The line in (b) was calculated similarly but used a 3-day rolling average. Dashed vertical lines indicate the dates of experimental trials.

### Response of heart rate and whole-animal thermal tolerance under acute warming

Critical thermal maximum (CT_max_), a measure of acute whole organism heat tolerance, was higher in summer, compared to winter (+4.5°C, emmeans seasonal contrast P<0.0001; Figure 3d; Table S2). The Acclimation Response Ratio (ARR; ΔCT_max_/Δaccl. body temp.) was 0.45 when calculated using winter and summer body temperature estimates. Relative ventricular mass was 19% greater in summer than winter (0.131 vs 0.110 % Body Mass; P=0.037). Overall, for both seasons and drug treatments, the *f*_H_ increased with acute temperature in all fish (emtrends, all 95% CIs above zero; P<0.0001; Figures 4a, b, Table S1). Drug treatment had little effect on **f**_H_ across most of the warming range in either season. However, *f*_H_ response to acute warming differed between seasons (χ^2^=48.3, P<0.0001), with *f*_H_ increasing to a greater extent in the summer than the winter.

**Figure 3.**
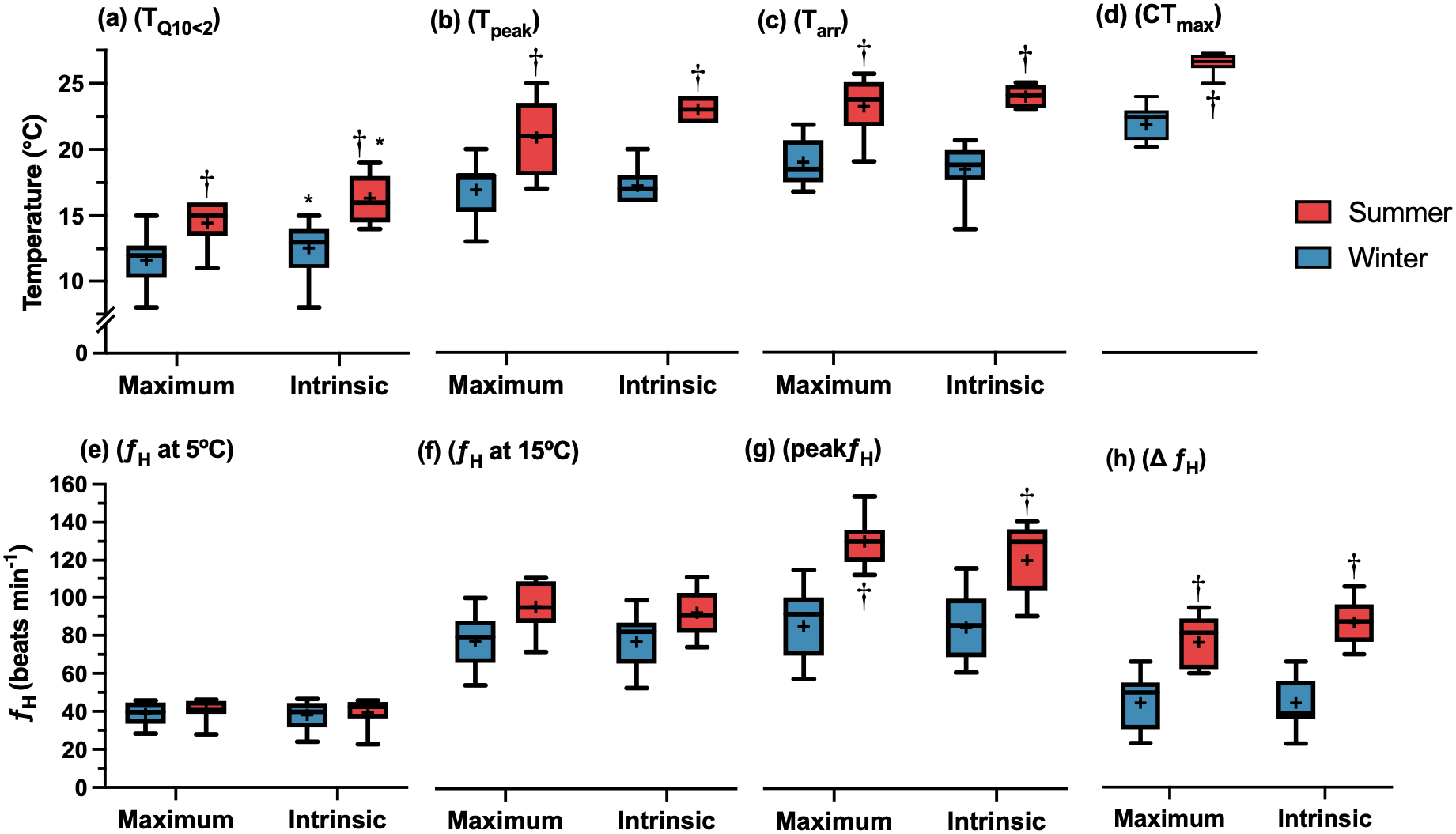
The effect of adrenergic stimulation (drug treatment) and seasonal acclimatization on heart rate (*f*_H_) and cardiac heat tolerance. (a) Temperature at which the incremental Q_10_ fell below 2, (b) temperature at peak heart rate, (c) temperature at arrhythmia, (d) temperature at critical thermal maximum, (e) heart rate at 5°C, (f) heart rate at 15°C, (g) peak heart rate, (h) change in heart rate between 5°C and peak heart rate. Boxes represent median and interquartile ranges, with whiskers as the 5^th^ and 95^th^ data percentile, crosses indicate the mean. Significant differences were found between seasons (P<0.01), except for heart rate at 5 and 15°C (e, f). Daggers (†) indicate differences between seasons within a drug treatment, and asterisks indicate differences between drug treatments within a season. Summer maximum n=9, summer intrinsic n=9, winter maximum n=12, winter intrinsic n=9, summer CT_max_ n =12, winter CT_max_ n = 11.

**Figure 4.**
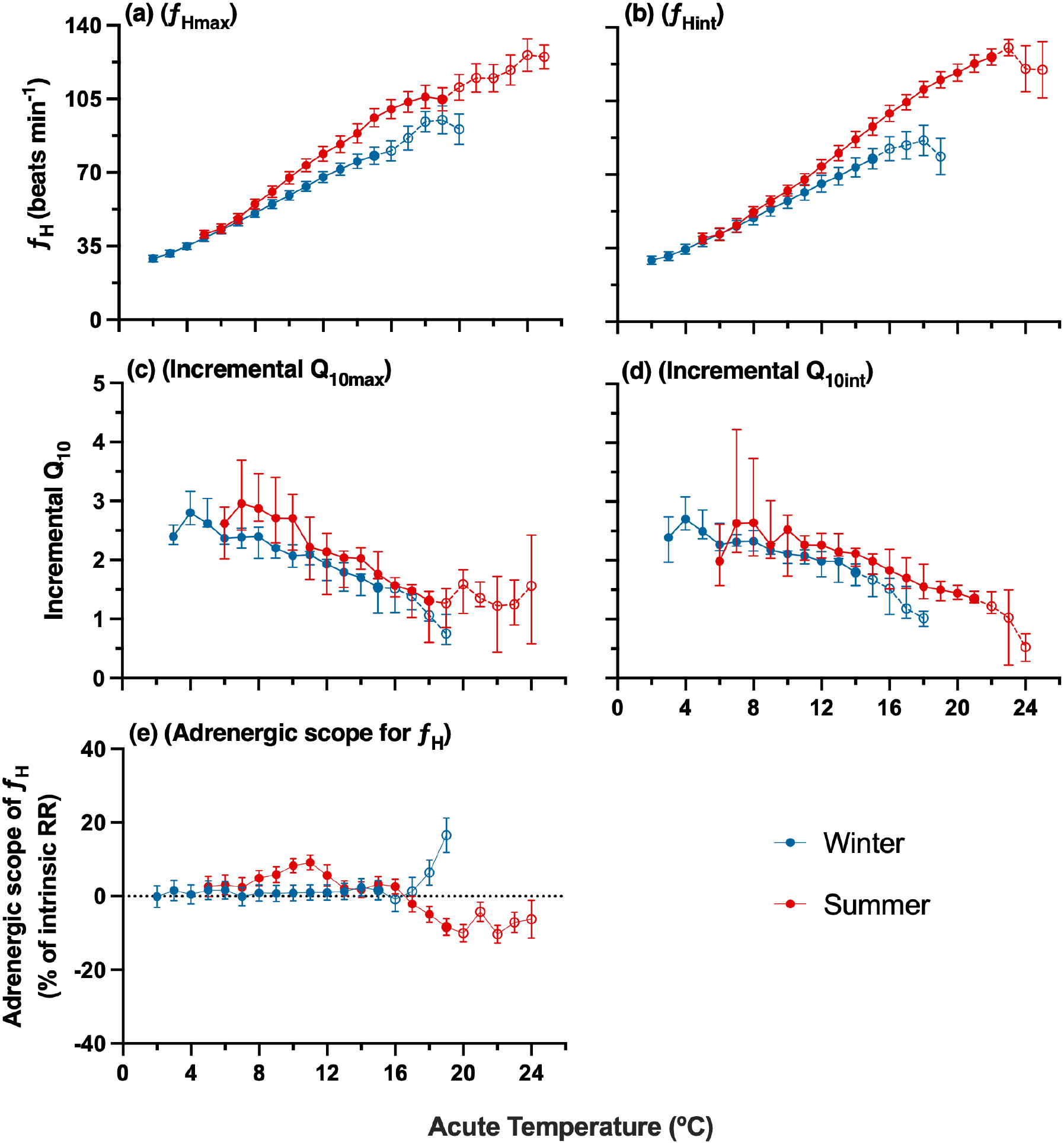
Acute warming effect on maximum (a) and intrinsic (b) heart rates and the associated incremental Q_10_ (c, d) and adrenergic scope of **f**_H_ (e) of seasonally acclimatized juvenile Arctic char. Estimated acclimatization temperatures (average body temperature during the experimental periods) were 1.1 °C in winter (n=21) and 11.1 °C in summer (n=18). Data presented as mean ± SEM with open circles and dashed lines indicating temperature at which hearts were arrhythmic.

Overall, thermal tolerance metrics differed between seasons, with summer-acclimatized fish generally exhibiting higher values than winter-acclimatized fish. Peak*f*_H_ differed strongly by season, with winter fish exhibiting peak heart rates approximately 40 BPM lower than summer fish (winter peak*f*_H_ 84.8±3.9, summer peak*f*_H_ 125.0±3.8 BPM, P<0.001; Figure 3g; Table S2). Drug treatment had no effect on peak heart rate (P=0.46). The Δ*f*_H_ was higher in summer compared to winter (P<0.0001), with no difference between drug treatments (Figure 3h; Table S2). Additional point comparisons were made at 5 and 15°C to account for summer fish not tested at 2°C (Figures 3e, f). Extrapolating **f**_H_ using model generated emmeans at 2°C found no projected difference at 2°C.

TQ_10_ (the temperature at which Q_10_ fell below 2) was higher in summer compared to winter-acclimatized fish (winter: 12.0±0.4°C, summer: 15.4±0.5°C, P<0.0001; Figure 3a; Table S2) and was higher in sotalol than isoproterenol-treated fish (P=0.03). T_peak_ also differed between seasons, with winter fish reaching peak heart rate at temperatures approximately 4.9°C lower than summer-acclimatized fish (winter: 17.0±0.5°C, summer: 21.9±0.4°C, ARR: 0.49, P<0.0001; Figure 3b; Table S2). We found weak evidence that drug treatment had a modest effect on T_peak_ (P=0.06), with sotalol-treated fish reaching peak heart rate at temperatures approximately 0.6°C higher than isoproterenol-treated fish. There was no interaction between season and drug treatment, suggesting that any effect of drug treatment on T_peak_ did not differ detectably between seasons. T_arr_ differed between seasons after accounting for body mass, with fish in winter starting to exhibit heat induced arrhythmia at 4.9°C lower than in summer (winter: 18.7±0.4°C, summer: 23.6±0.4°C, ARR: 0.49, P=0.001; Figure 3c; Table S2). Drug treatment had no effect on T_arr_ (P=0.93). Adrenergic scope for heart rate showed limited differences seasonally, with a slight decline apparent in summer acclimatized fish at higher temperatures (Figure 4e).

Incremental Q_10_ declined during acute warming in all fish over both seasons and drug treatments (emtrends at mean body mass, all 95% CIs are below zero; Figures 4c, d; Table S1). The rate of decline depended on season, drug treatment, and body mass (χ^2^=22.6, P<0.0001). Under isoproterenol treatment, winter-acclimatized fish exhibited a steeper decline in incremental Q10 with increasing temperature than summer-acclimatized fish (emtrends summer – winter slope contrast = 0.031±0.008, z=4.06, P<0.0001).

The PR interval decreased with acute warming in all fish across seasons and drug treatments (all 95% CI’s below zero; Figures 5a,b; Table S3). The PR interval was longer in winter than summer and the slope of decrease during warming depended on season (χ^2^=48.5, P<0.0001), drug treatment (χ^2^=41.1, P<0.0001) and body mass (χ^2^=104.8, P<0.0001). At low temperatures PR interval was initially shorter with β-adrenergic stimulation (isoproterenol) than without (sotalol), but the two treatments converged with warming (Figure S4; Table S3).

**Figure 5.**
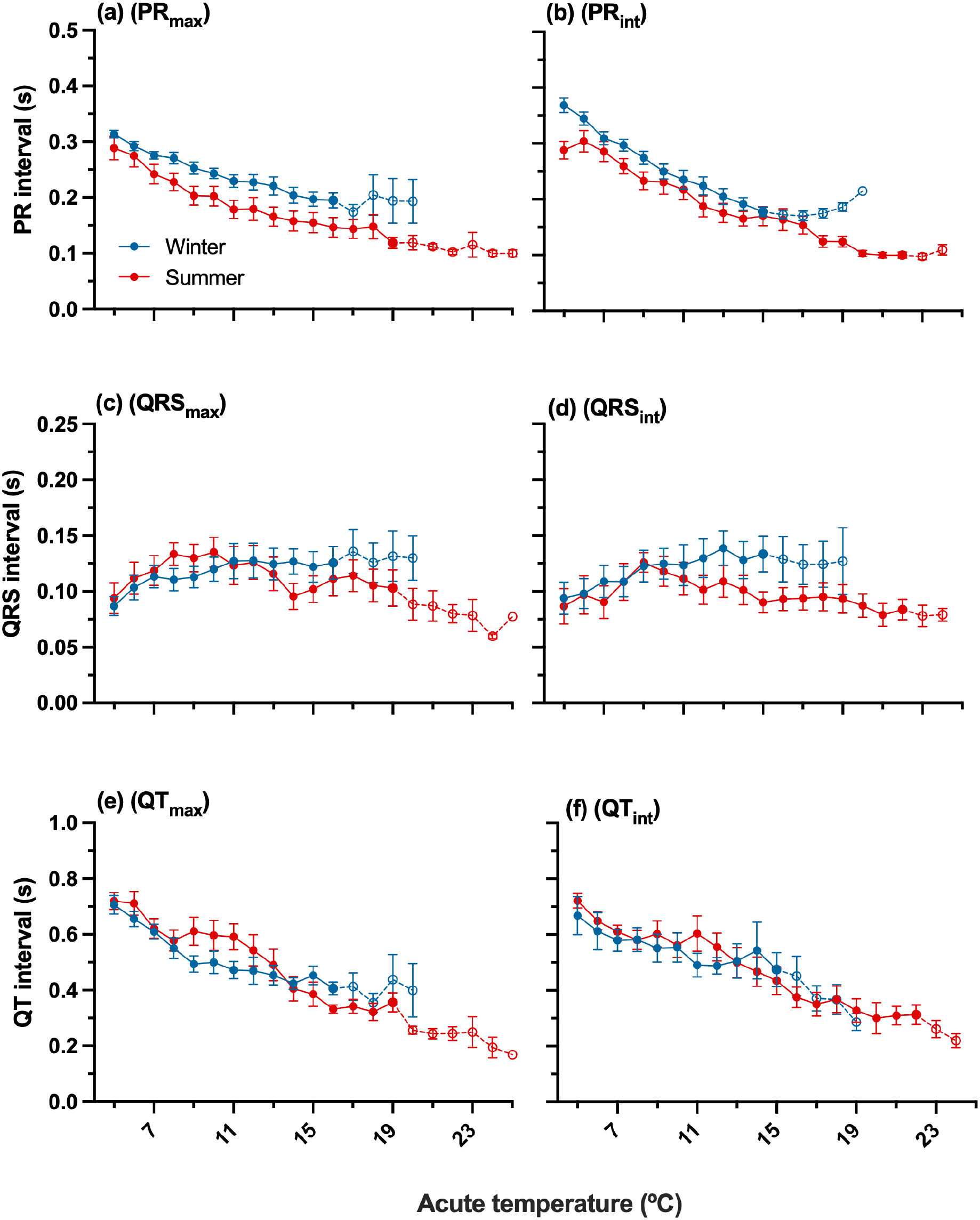
Effect of seasonal acclimatization and acute warming on the PR interval (a, b), QRS duration (c, d), and QT interval (e, f) on maximum (a, c, e) and intrinsic (b, d, f) heart rate. Data are presented as mean ± SEM, open circles and dashed lines indicate where fish were removed after onset of arrhythmia. Estimated acclimatization temperatures (average body temperature during the experimental periods) were 1.1 °C in winter (n=21) and 11.1 °C in summer (n=18).

The QT interval decreased with increasing temperature in all fish across both seasons and drug treatments (emtrends, all 95% CIs below zero; Figures 5e, f; Table S4). At 5°C, QT intervals did not differ between seasons (summer/winter ratio = 1.01 ± 0.09, z = 0.12, *P* = 0.91) or drug treatments (ISO/SOT ratio = 0.99 ± 0.08, z = −0.16, *P* = 0.87). The temperature dependence of the QT interval differed between seasons (χ²=5.7, *P*=0.02), with QT decreasing more steeply with warming in summer-than in winter-acclimatized fish (summer − winter slope contrast = −0.01 ±0.005, z=-2.4, P=0.02). QT also decreased more steeply under adrenergic stimulation than under adrenergic blockade (ISO − SOT slope contrast = −0.011±0.004, z =-2.9, P = 0.004; χ^2^=8.3). In both summer and winter fish, diastolic duration for *f*_Hmax_ and *f*_Hint_ decreased with acute warming (Figures S2a, b; Table S5). The ratio between diastolic and systolic duration was seen to increase in summer fish once ∼16°C was reached for both drug treatments, indicating an increase in the duration of ventricular diastole (Figures S2c, d). QRS duration responded differently to acute warming between seasons (χ^2^ = 153.0, P < 0.0001), increasing with temperature in winter-acclimatized fish (slope=0.004±0.0003) and decreasing with warming in summer-acclimatized fish (slope= −0.002±0.0003; Figures 5c, d, Table S3). Drug treatment had no effect on QRS duration (χ^2^=0.2, P=0.67).

### Cardiac arrhythmia analysis

Juvenile Arctic char experienced several types of cardiac arrhythmias during acute warming. In winter, five fish treated with isoproterenol experienced asystole, as did six fish treated with sotalol (Figures 6c.1, 2). Three isoproterenol-treated fish experienced type 1 second-degree AV-block where the PR-interval was prolonged. Four isoproterenol-treated fish experienced type 2 second-degree AV block with a dissociation of the p-wave and QRS complex. Two sotalol-treated fish experienced type 1 second-degree AV block, and one experienced type 2 second-degree AV block. In the summer, three isoproterenol-treated fish experienced asystole, as did seven sotalol-treated fish (Figures 6d.1, 2). Six isoproterenol-treated fish experienced AV block, with two showing type 1 second-degree, and four showing type 2 second-degree. One sotalol-treated fish experienced type 1 second-degree AV block, and one experienced type 2 second-degree.

**Figure 6.**
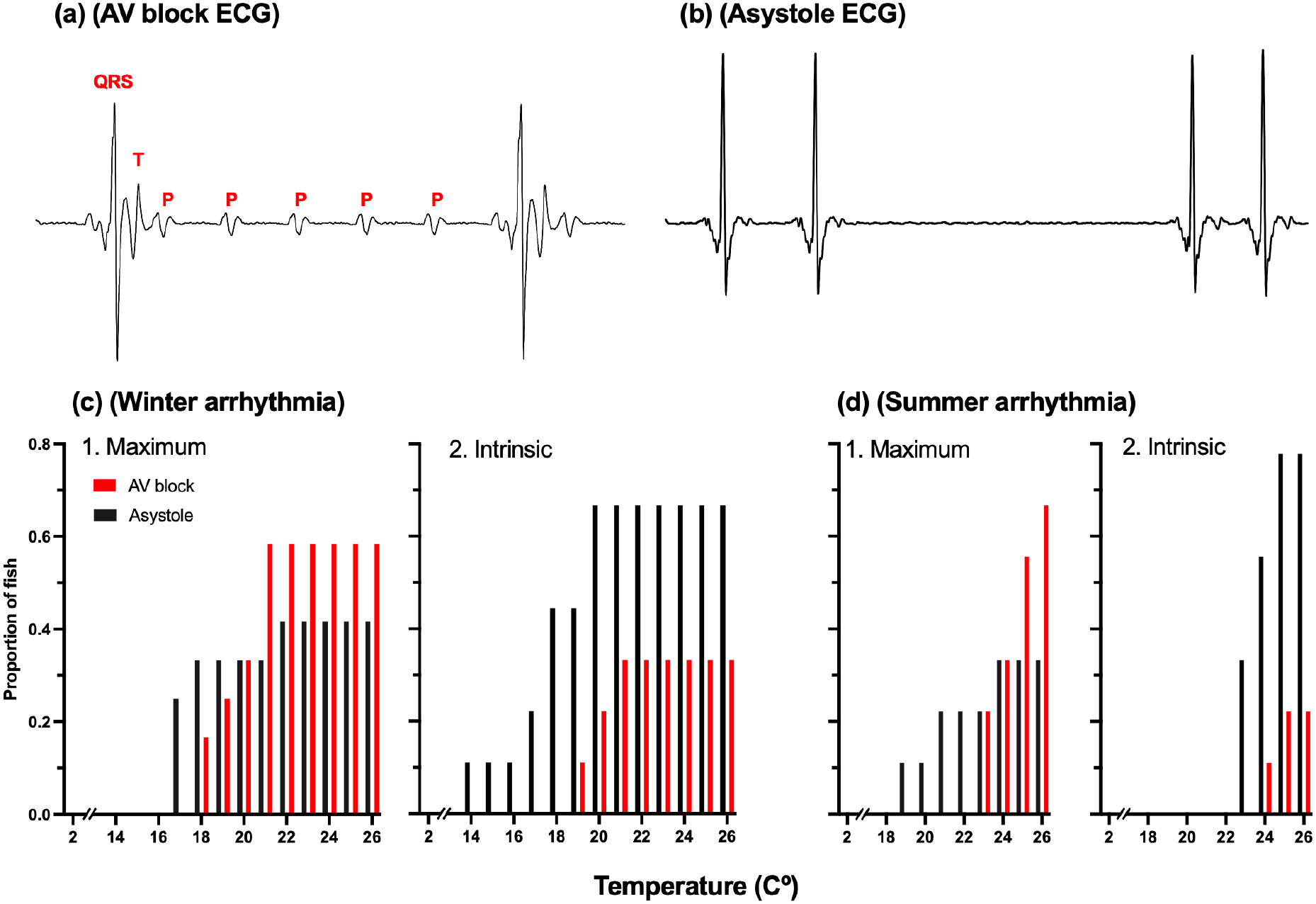
Juvenile Arctic char electrocardiograph tracings showing a section of AV block (a) of a summer acclimatized fish at 21 °C, and a section of asystole (b) of a summer acclimatized fish at 19 °C. Frequency histograms for winter (c) and summer (d) proportion of acclimatized Arctic char experiencing either AV block or asystole-type arrhythmias, separated by drug treatment of maximum; isoproterenol (1) and intrinsic; sotalol (2).

## Discussion

High-latitude Arctic char can encounter substantial thermal variation with temperatures ranging from <1°C in winter to >17°C during summer in freshwater lakes, and even higher temperatures during summer upriver migrations (Gilbert *et al*., 2016). To maintain critical physiological functions across such thermal extremes, many fishes exhibit thermal compensation of biological rates and thermal tolerance as a result of thermal acclimation (Haverinen and Vornanen, 2007; Vornanen *et al*., 2014; Gilbert *et al*., 2020; Gilbert and Farrell, 2021). Here, we found that average body temperatures increased from 1.2°C in winter to 8.9°C in summer, providing a strong potential for such thermal plasticity to occur. Accordingly, juvenile Arctic char exhibited evidence of seasonal acclimatization in CT_max_, the responsiveness of *f*_H_ to warming, T_peak_, and T_arr_. Seasonal acclimatization also affected the electrical activity of the heart, with the PR interval being prolonged in winter-acclimatized fish, and QRS being longer at warmer temperatures in winter, but with QT showing limited differences. Corroborating our hypothesis, we found supporting evidence that juvenile Arctic char demonstrate strong seasonal plasticity in cardiac and whole-body thermal performance, processes that have been unquantified seasonally for juvenile Arctic char. In contrast to our hypothesis and laboratory thermal acclimation studies, we did not find evidence of compensatory pacemaker resetting. We also observed limited seasonal differences in adrenergic scope for heart rate. Overall, although certain aspects of cardiac function of wild juvenile Arctic char showed seasonal acclimatization, others differed from what has been observed in lab and in wild adult Arctic char. Thus, this study illustrates the importance of life-stage-specific and field-based research.

### Whole animal and cardiac heat tolerance

CT_max_ remains the most widespread indicator of the absolute upper bounds for whole animal thermal performance and is often measured using similar methodology allowing for comparisons between and within fish species across studies (Sandblom *et al*., 2016). However, the direct ecological relevance of CT_max_ measurements can be limited. In the wild, such rapid and extreme high temperatures are rarely within a fish’s range and fish will experience other severe functional limitations before reaching CT_max_ (Desforges *et al*., 2023). Previous studies of hatchery-raised European Arctic char juveniles (20-50 g) found small change in CT_max_ with thermal acclimation across intermediate temperatures. Juveniles acclimated to 7.7 and 14.9°C showed only a 1.3°C increase in CT_max_, whereas fish acclimated to 5 and 10°C showed a 0.5°C increase (Baroudy and Elliott, 1994; Anttila *et al*., 2015). In larger, intermediately sized lab-raised Arctic char (400 g) acclimated to 2 and 10°C, CT_max_ increased from 24.5 to 26.5°C, with an ARR of 0.23 (Gilbert and Farrell, 2021). Wild temperate brook trout acclimatized to a 30-day temperature mean of 9.09°C and 17.90°C increased CT_max_ from 28.09 to 30.09°C, resulting in an ARR of 0.23 as well (Stewart *et al*., 2024). These differences were smaller than the seasonal difference observed in the present study, where CT_max_ increased by 4.5°C (from 21.4 to 25.9°C) between winter and summer, with a ∼10°C increase in acclimatization temperature and an ARR of 0.45. In summer, CT_max_ was approximately 3°C higher in the juveniles examined here than in larger migrating Arctic char (∼500-920 g) caught in a prior study in a nearby river at water temperatures ranging from 8.9 to 14.5°C (CT_max_: 23.0°C; Gilbert *et al*., 2020) and in hatchery-raised adult Arctic char acclimated to ∼10°C (CT_max_: 23.0°C; Penney *et al*., 2014). While the juveniles in this study show substantial seasonal plasticity in their acute critical whole-animal heat tolerance and high summer CT_max_, it is essential to consider that fish may experience important physiological limitations including impaired cardiorespiratory performance at temperatures below their CT_max_ (Chen *et al*., 2013; Gilbert *et al*., 2020, 2024; Gilbert and Farrell, 2021).

Acute thermal limitations such as cardiac arrhythmia and heart rate collapse typically occur at temperatures below CT_max_. Because fish are more likely to encounter these lower temperatures in the wild, such limits may be more ecologically relevant than CT_max_ itself. For instance, Gilbert *et al*. (2020) found that the distribution of T_peak_ (19.4 ± 0.5°C) and T_arr_ (21.4 ± 0.5°C) in upstream migrating adult Arctic char had substantial overlap with upper river temperatures during heat waves in which water reached 21°C. Likewise, Mottola *et al*. (2020) found that T_arr_ was only 2°C above typical migratory river temperatures. However, neither of these studies examine the ability of these fish to increase heat tolerance through acclimatization. In a laboratory study, Gilbert and Farrell (2021) found that T_arr,_ T_peak_, and TQ_10_ increased by 35%, 49%, and 37%, respectively, with acclimation from 2 to 14°C. When comparing fish acclimated to 2 and 10°C, the same study reported ARR values of 0.54 for T_arr_ and 0.58 for T_peak_. A field-experiment found that these changes can begin within 72hr (Gilbert *et al*., 2022b). Our findings further provide evidence that Arctic char have capacity for cardiac acclimatization that may help buffer against high temperature events. We saw an increase in T_arr_, T_peak_, and TQ_10_ from 18.7 ± 0.4°C, 17.0 ± 0.5°C, and 12.0 ± 0.4°C in winter acclimatized juveniles to 23.6 ±0.4°C (ARR: 0.49), 21.9 ±0.4°C (ARR: 0.49), and 15.4 ± 0.5 °C (ARR: 0.34) in summer acclimatized juveniles. Lastly, the wild juveniles in this study showed no obvious protective effect of adrenaline on these thermal tolerance metrics. Similarly, Ekström *et al*. (2014) found no effect of adrenaline on CT_max_ in rainbow trout *Oncorhynchus mykiss*, a temperate salmonid. However, in a similar study using comparable methods, adrenaline was found to enhance acute cardiac heat tolerance in rainbow trout (Gilbert *et al*., 2019). Thus, the role of adrenaline in heat tolerance can differ even within species in similar contexts. Such differences may arise because autonomic cardiac control is plastic, varying with life stage, thermal history, and migration intensity.

### Maximum and intrinsic heart rate

Resetting cardiac pacemaker rates through modification of sinoatrial (SA) currents is one mechanism many fish use to compensate for sustained periods of temperature change to offset the depressive effects of cooling and stimulatory effects of warming (Haverinen and Vornanen, 2007; Anttila *et al*., 2014; Badr *et al*., 2016; Eliason and Anttila, 2017; Sutcliffe *et al*., 2020). This response has been observed in several species (e.g. Roach: *Rutilus rutilus;* rainbow trout: *Oncorhynchus mykiss*) (Aho and Vornanen, 2001; Anttila *et al*., 2014; Gilbert *et al*., 2022a), including in Arctic char which showed a reduction in heart rate following warm acclimation in lab- and field-based experiments (Gilbert and Farrell, 2021; Gilbert *et al*., 2022b). In contrast, here, wild juvenile Arctic char showed no such resetting of initial *f*_H_ values (when compared at 5°C). Likewise, Williams *et al*. (2025), found nearly identical *f*_H_ values for cold and warm-acclimatized individuals over intermediate temperatures which highlights that this response is not universal and can be specific to the species, life stage, study setting, or temperature range the fish is measured at. The lack of *f*_H_ compensation (i.e., no resetting response) in summer acclimatized fish could indicate a greater reliance on stroke volume to maintain cardiac output, or a reduced demand for cardiac output due to decreased oxygen demand after cooling. The higher RVM seen in summer-acclimatized fish is consistent with supporting a greater demand of stroke volume. Alternatively, a compensatory increase in **f**_H_ may have occurred but been offset by the overall depressive effect of the cold on metabolic rate, including that of the heart (Middleton *et al*., 2024; Gilbert *et al*., 2026). While we did not see resetting of pacemaker rates, it is important to note that we could not directly compare heart rate at temperatures below 5°C because summer acclimatized fish exhibited cold-induced arrhythmias during preliminary trials when cooled to 2°C. Likewise, Gilbert *et al*. (2020) found that wild upriver migrating Arctic char displayed cold-induced arrhythmias, with a collapse in heart rate seen below 3.3°C (Gilbert *et al*., 2020). Thus, while pacemaker rates did not change, summer acclimatization appears to compromise stability of cardiac electrical activity when cooled. Interestingly this arrhythmia did not occur in the lab intermediate-sized char or in a separate study of wild ocean caught adult Arctic char (Gilbert and Farrell, 2021; Gilbert *et al*., 2022b), again demonstrating that acclimation and acclimatization response can be highly context dependent.

Beyond resetting of *f*_H_, β-adrenergic control of cardiac function can change with thermal acclimation and seasonal acclimatization (Williams et al., 2025). Such β-adrenergic control can, but does not always, enhance aspects of cardiac and whole-animal heat tolerance (Ekström *et al*., 2014; Gilbert *et al*., 2019). Gilbert *et al*. (2019) found that adrenergic stimulation increased cardiac and whole-animal heat tolerance, with β-adr (β-adrenergic) blockade lowering peak*f*_H_ by 14-17% and CT_max_ by 3°C, while Ekström *et al*. (2014) did not find similar benefits of β-adrenergic stimulation. In a field study that overlapped with the present study (same methods and sampling locations) Williams *et al*. (2025) found that peak*f*_H_ during acute warming was lower under β-adr blockade, however this difference only occurred in winter, and was not accompanied by any difference in T_peak_ or T_arr_. Williams et al (2025) also found a β-adr**f**_H_ scope of ∼15-30% over sub-critical temperatures in alignment with several other studies in salmonids using varied methodologies (Ekström et al., 2016; Gilbert et al., 2019). Here, we found no effect of β-adr blockade on peak*f*_H_, T_arr,_ or T_peak_, with only TQ_10_ having higher values, and surprisingly found that sotalol had little effect on heart rate across most of the warming range in either season (limited β-adr**f**_H_ scope). Thus, while we did see minor seasonal changes in β-adr control of cardiac electrical activity (see section below) they do not appear to underlie observed seasonal differences in cardiac heat tolerance.

### Cardiac arrhythmia and electrical activity

During acute warming, the PR interval of juvenile Arctic char decreased regardless of season and drug treatment, indicating that atrioventricular (AV) conduction rates increased with warming and heart rate, as expected. However, even at similar heart rates the PR interval was longer in winter than summer. Further PR interval appeared to be shortened by β-adrenergic stimulation at cold temperatures, with the effect being diminished as warming progressed. Because cholinergic receptors were blocked with atropine, the longer PR interval in winter is unlikely to reflect differences in vagal tone and instead suggests intrinsic remodeling of AV conduction. The PR interval depends largely on the duration of the AV nodal delay and the rate of cardiac action potential (AP) upstroke, which is mainly determined by the density of the inward sodium current (I_Na_), and of the calcium current (I_Ca_) in nodal tissue (Vornanen, 2017). In atrial myocytes, density of I_Na_ tends to be larger than I_Ca_ so the duration of the atrial conduction is largely influenced by I_Na_ (Haverinen and Vornanen, 2006). Thus, a longer PR interval in winter-acclimatized fish could indicate that I_Na_ density is depressed with cold acclimatization as previously found in crucian carp (*Carassius carassius*) and European shorthorn sculpin (*Myoxocephalus scorpio*) (Haverinen and Vornanen, 2004; Filatova *et al*., 2019). However, the direction of INa remodeling with cold appears to be species-or context-specific, so it may not be the main driver of reduced cardiac heat tolerance. For example, rainbow trout have been found to increase I_Na_ density with cold acclimation (Haverinen and Vornanen, 2004) but commonly show similar reductions in cardiac heat tolerance with reductions in acclimation temperature (Gilbert *et al*., 2022a; Strowbridge *et al*., 2025).

Potassium (K^+^) currents, specifically the delayed rectifier current (I_Kr_) have also been shown to be upregulated with cold acclimation, and can play an important role in seasonal acclimation of cardiac excitability (Hassinen *et al*., 2008; Abramochkin and Vornanen, 2015; Vornanen, 2017; Filatova *et al*., 2019). Greater K^+^ current density shortens action potential duration thereby decreasing QT interval (Vornanen, 2017). Here, juvenile Arctic char showed minimal seasonal differences in QT interval, which may indicate a limited seasonal remodeling of repolarizing K⁺ currents, unlike what is commonly observed with laboratory cold acclimation in other species. The steeper decline in QT interval with warming in summer fish may instead reflect their higher heart rates at warm temperatures, as QT shortens with increasing heart rate. In contrast, QRS durations at intermediate to high temperatures tended to be longer in winter, as shown in lake trout and roach, which is consistent with slower ventricular depolarization rates (Badr *et al*., 2016; Williams *et al*., 2025). However, unlike those studies in which QRS duration declined consistently during acute warming, juvenile Arctic char in both seasons showed an initial increase in QRS duration up to ∼10°C, the cause of which is uncertain, before plateauing in winter fish and decreasing in summer-acclimatized fish. The seasonal divergence at higher temperatures, where QRS duration remained long in winter fish as they approached their upper thermal limits, suggests that a ventricular conduction limitation, possibly arising from a source–sink mismatch (Vornanen, 2020), may contribute to their lower cardiac heat tolerance. Impulse transmission requires that the depolarizing source current (I_Na_) exceeds the repolarizing sink current that maintains the resting potential of receiving cells (I_K1_; outward K^+^ current, Vornanen, 2020). During the initial rise in temperature, and associated increase in heart rate, overall shortened diastolic intervals, as seen in juvenile char, could also reduce recovery time for Na⁺ channels, which recover slowly from inactivation at low temperatures, potentially slowing excitability and conduction and contributing to the temporary increase in QRS duration. Additionally, a depressed I_Na_ density in winter could contribute to AV block occurring at lower temperatures in winter and the reduction in cardiac heat tolerance. The large seasonal differences observed may therefore arise from a source-sink mismatch as well rather than changes in β-adrenergic sensitivity. In fish hearts, I_Na_ is heat-sensitive and declines with warming, whereas I_K1_ is heat-resistant and increases with rising temperatures (Vornanen *et al*., 2014; Badr *et al*., 2018), so the source decreases as the sink increases until conduction fails between the atria and ventricle. Winter-acclimatized, isoproterenol-treated fish also tended to experience AV block-type arrhythmias more than sotalol-treated fish (7 of 12 vs. 3 of 9), in which atrial impulses fail to be conducted through the AV node to the ventricle (Vornanen, 2020). However, sample sizes were small and β-adrenergic stimulation shortened rather than prolonged the PR interval at cold temperatures, so the role of adrenergic stimulation in AV block remains unclear. Combined, although some seasonal differences in ECG waveforms were small, they suggest that seasonal electrical remodeling of the heart occurs in juvenile Arctic char in a manner that likely influences cardiac thermal sensitivity.

### Ecological implications

Juvenile Arctic char express marked whole-animal and cardiac seasonal plasticity, that aligns with their persistence in highly variable Arctic freshwater environments. However, with the Arctic warming at a rapid pace (Rantanen *et al*., 2022), this may impair cardiac function thereby constraining whole-animal performance. In 2023, the region experienced a remarkably warm summer, with temperatures in a neighboring Arctic char-bearing lake (Greiner Lake) reaching 18°C (Williams *et al*., 2025). Although the maximum temperature we recorded in Inuhuktok was 13.5°C, the lake experiences substantial seasonal temperature variation and could experience higher summer temperatures with continued warming. While summer-acclimatized juveniles in this study had T_arr_ and T_peak_ above 18°C, the intermediately-sized lab acclimated char in Gilbert and Farrell (2021) exhibited increased mortality and the cessation of feeding following prolonged exposure (5-7 weeks) to 18°C. Thus, if juvenile Arctic char in this watershed encounter prolonged periods at such temperatures, growth, survival, and habitat use could be affected, potentially influencing population dynamics and broader ecosystem interactions. Many Arctic lakes are thought to not thermally stratify, which can limit the availability of cold-water refugia during heat events for both present and future landlocked char. This narrative is changing as the prevalence of stratification at high latitudes has been recently documented in Arctic lakes (Antoniades *et al*., 2024) and indeed Inuhuktok (present study lake) stratified in the summer of 2026 for the first time since monitoring began in 2018 (unpublished data). Additionally, as seen during the heatwave that occurred in 2023, neighboring lakes can experience different thermal regimes, which may have important implications for habitat connectivity because many fish in this system move amongst lakes. If all lakes have warmed and one becomes markedly hotter than others, fish may avoid it, resulting in an effective loss of habitat. For example, juvenile char in the region, including those from Inuhuktok, must pass through Greiner Lake to access a creek used as foraging habitat. If Greiner Lake, and likely the creek itself, experience higher temperatures than other lakes in the region, fish may reduce their use of these habitats, alter the timing of movements, or may only be able to utilize them for short times.

Here, we have shown that wild juvenile Arctic char exhibit notable seasonal plasticity by modulating the temperature sensitivity of both whole-animal and cardiac performance. However, because many responses to temperature and season are species- and context-dependent, thermal limits studied under a limited range of conditions may be inaccurately estimated. For instance, as juvenile Arctic char show acute sublethal limitations well below CT_max_, future studies should examine additional functional thermal limitations over more ecologically relevant and prolonged timescales than shown here. Such data can provide Indigenous Rightsholders and managers with information that, when integrated with environmental data and predictive models, can help identify areas of concern and predict responses to changing environments, such as the likelihood of mortality events or changes in growth patterns. Thermal performance data can also help inform the identification of emerging conservation concerns and support decision-making within integrated fisheries management plans (DFO, 2021). Together, these approaches place acute thermal performance measurements within a broader understanding of the physiological and ecological responses of Arctic char to environmental change.

## Acknowledgements

Quana (thank you) to the community of Ikaluktutiak, Cambridge Bay, Nunavut and to the Ekaluktutiak Hunters and Trappers Organization (EHTO) for their support of this research. The authors thank Kevin Kanayok, Richard Ekpakohak, Vincent Panaktak, Brent Nakashook, and Elise Imbeau for contributions as field technicians and with sampling help. Thank you to Kristin O’Brien and Erik Schoen for valuable feedback that improved the clarity of this manuscript. This research was supported by Polar Knowledge Canada. SAB was supported by the Calvin J. Lensink Graduate Fellowship and the Jack R. Luick Memorial Travel Endowment. Field sampling and animal use were approved by the UAF Institutional Animal Care and Use Committee ( # 2181369-3, # 2302455-1) and Fisheries and Oceans Canada (OPA ACC-2024-40, OPA-ACC-2025-3).

## Notes

### Competing Interest Statement

The authors have declared no competing interest.

