## Supplemental Material for "Wild juvenile Arctic char (*Salvelinus alpinus*) show seasonal acclimatization of cardiac and whole-animal heat tolerance"

**Table S1**. Results of the generalized linear mixed-effect models examining the change in heart rate ($f_H$) and incremental $Q_{10}$ under acute warming for maximum and intrinsic heart rate treatments and seasonal treatments (winter and summer) of wild juvenile Arctic char. The $f_H$ was modeled from 5°C to peak$f_H$ and $Q_{10}$ from 6°C to peak$f_H$. Fish ID was assigned as the random effect. Significant effects ($P < 0.05$) calculated using a type III Wald chi-square test.

| **Fixed effect** | **Chi-square** | **df** | ***P*** |
|---|---|---|---|
| ***$f_H$* (beats min$^{-1}$)** | | | |
| Intercept | 325.1 | 1 | $< 2.2 \times 10^{-16}$ |
| Body mass (g) | 0.1 | 1 | 0.74 |
| Acute temperature (°C) | 1739.1 | 1 | $< 2.2 \times 10^{-16}$ |
| Season | 6.1 | 1 | 0.01 |
| Drug treatment | 1.8 | 1 | 0.17 |
| Acute temperature x Season | 48.3 | 1 | $3.6 \times 10^{-12}$ |
| Acute temperature x Drug | 23.0 | 1 | $1.6 \times 10^{-6}$ |
| Season x Drug | 0.8 | 1 | 0.36 |
| Acute temperature x Season x Drug | 12.1 | 1 | $5.17 \times 10^{-4}$ |
| **Incremental $Q_{10}$** | | | |
| Intercept | 16.4 | 1 | $5.24 \times 10^{-5}$ |
| Body mass (g) | 20.5 | 1 | $5.95 \times 10^{-6}$ |
| Acute temperature (°C) | 3.4 | 1 | 0.06 |
| Season | 17.8 | 1 | $2.45 \times 10^{-5}$ |
| Drug treatment | 16.1 | 1 | $5.94 \times 10^{-5}$ |
| Mass x Temperature | 55.8 | 1 | $8.18 \times 10^{-14}$ |
| Mass x Season | 18.9 | 1 | $1.37 \times 10^{-5}$ |
| Temperature x Season | 86.1 | 1 | $< 2.2 \times 10^{-16}$ |
| Mass x Drug | 12.0 | 1 | $5.28 \times 10^{-4}$ |
| Temperature x Drug | 69.7 | 1 | $< 2.2 \times 10^{-16}$ |
| Season x Drug | 13.3 | 1 | $2.68 \times 10^{-4}$ |
| Mass x Temperature x Season | 87.6 | 1 | $< 2.2 \times 10^{-16}$ |
| Mass x Temperature x Drug | 67.6 | 1 | $< 2.2 \times 10^{-16}$ |
| Mass x Season x Drug | 6.6 | 1 | 0.01 |
| Temperature x Season x Drug | 44.9 | 1 | $2.10 \times 10^{-11}$ |
| Mass x Temperature x Season x Drug | 22.6 | 1 | $2.00 \times 10^{-6}$ |

Model: glmer(HR~Mass+Temperature*Season*Drug +(1|ID), ARCH, family=Gamma(link='log'))
Model: glmer(IncQ10~Mass*Temperature*Season*Drug +(1|ID), ARCH, family=Gamma(link='log'))

**Table S2**. Results of the linear models statistics for the change in $T_{peak}$ (temperature at peak heart rate), Peak$f_H$ (peak heart rate), change in heart rate between 5ºC and peak heart rate, $T_{arr}$ (temperature at arrhythmia), $TQ_{10}$ (temperature at which the incremental $Q_{10}$ fell below 2), and $CT_{max}$ (temperature at critical thermal maximum) under acute warming for maximum and intrinsic heart rate treatments and seasonal treatments (winter and summer) of wild juvenile Arctic char.

| Fixed effect | Sum of squares | df | *F* value | *P* |
|---|---|---|---|---|
| **$T_{Q10}$** | | | | |
| Intercept | 7310.1 | 1 | 2096.7 | $< 2.2 \times 10^{-16}$ |
| Season | 101.5 | 1 | 29.1 | $4.46 \times 10^{-6}$ |
| Drug treatment | 17.7 | 1 | 5.1 | 0.03 |
| **$T_{peak}$ (°C)** | | | | |
| Intercept | 14612.0 | 1 | 3893.2 | $< 2.2 \times 10^{-16}$ |
| Season | 228.2 | 1 | 60.8 | $3.69 \times 10^{-9}$ |
| Drug treatment | 14.0 | 1 | 3.7 | 0.06 |
| Season x drug treatment | 7.8 | 1 | 2.1 | 0.16 |
| **Peak $f_H$ (beats min$^{-1}$)** | | | | |
| Intercept | 425454.0 | 1 | 1431.2 | $< 2.2 \times 10^{-16}$ |
| Season | 15266.0 | 1 | 51.4 | $1.99 \times 10^{-8}$ |
| Drug treatment | 169.0 | 1 | 0.6 | 0.46 |
| **$\Delta f_H$ (beats min$^{-1}$)** | | | | |
| Intercept | 166526.0 | 1 | 826.8 | $< 2.2 \times 10^{-16}$ |
| Season | 14115.0 | 1 | 70.1 | $5.72 \times 10^{-10}$ |
| Drug treatment | 248.0 | 1 | 1.2 | 0.27 |
| **$T_{arr}$ (°C)** | | | | |
| Intercept | 622.6 | 1 | 214.7 | $< 2.2 \times 10^{-16}$ |
| Body mass (g) | 10.5 | 1 | 3.6 | 0.07 |
| Season | 37.3 | 1 | 12.9 | $1.01 \times 10^{-3}$ |
| Drug treatment | 0.0 | 1 | $8.1 \times 10^{-3}$ | 0.93 |
| **$CT_{max}$** | | | | |
| Intercept | 342.3 | 1 | 467.17 | $7.73 \times 10^{-15}$ |
| Season | 0.09 | 1 | 0.12 | 0.73 |
| Body mass (g) | 4.3 | 1 | 5.80 | 0.03 |
| Season x body mass | 3.7 | 1 | 5.05 | 0.04 |

Model: lm(TQ10 ~ Season + Drug, TQ10)
Model: lm(Tpeak ~ Season*Drug, ARCHTol)
Model: lm(Peakfh ~ Season + Drug, ARCHTol)
Model: lm(Delta_fh ~ Season + Drug, ARCHTol_delta)
Model: lm(Tarr ~ BodyMass + Season + Drug, ARCHTol)
Model: lm(Ctmax ~ Season*BodyMass, ARCH_ctmax_2)

**Table S3.** Results of the generalized linear mixed effect model (PR interval) and linear mixed effect model (QRS duration) statistics for the change in PR interval and QRS duration under acute warming for maximum and intrinsic heart rate treatments and seasonal treatments (winter and summer) of wild juvenile Arctic char. Both PR interval and QRS duration were modeled from 5°C to peak $f_H$. Fish ID was assigned as the random effect. Significant effects ($P < 0.05$) calculated using a type III Wald chi-square test.

| **Fixed effect** | **Chi-square** | **df** | ***P*** |
|---|---|---|---|
| **PR interval (s)** | | | |
| Intercept | 42.1 | 1 | $8.46 \times 10^{-11}$ |
| Body mass (g) | 0.2 | 1 | 0.64 |
| Acute temperature (°C) | 1221.7 | 1 | $< 2.2 \times 10^{-16}$ |
| Season | 1.4 | 1 | 0.23 |
| Drug treatment | 13.7 | 1 | $2.20 \times 10^{-4}$ |
| Mass x Temperature | 104.8 | 1 | $< 2.2 \times 10^{-16}$ |
| Temperature x Season | 48.5 | 1 | $3.24 \times 10^{-12}$ |
| Temperature x Drug | 41.1 | 1 | $1.46 \times 10^{-10}$ |
| **QRS duration (s)** | | | |
| Intercept | 357.2 | 1 | $< 2.2 \times 10^{-16}$ |
| Acute temperature (°C) | 19.7 | 1 | $8.84 \times 10^{-6}$ |
| Season | 24.6 | 1 | $7.10 \times 10^{-7}$ |
| Drug treatment | 0.2 | 1 | 0.67 |
| Temperature x Season | 153.0 | 1 | $< 2.2 \times 10^{-16}$ |

Model: glmer(PR ~ Mass*Temperature + Temperature*Season + Temperature*Drug + (1|ID), ARCH, family=Gamma(link='log'))
Model: lmer(QRS ~ Temperature * Season + Drug + (1|ID), data = ARCH)

**Table S4.** Results of the generalized linear mixed effect models for the change in RR and QT interval under acute warming for maximum and intrinsic heart rate treatments and seasonal treatments (winter and summer) of wild juvenile Arctic char. Both RR and QT intervals were modeled from 5°C to peak$f_H$. Fish ID was assigned as the random effect. Significant effects ($P < 0.05$) calculated using a type III Wald chi-square test.

| **Fixed effect** | **Chi-square** | **df** | ***P*** |
|---|---|---|---|
| **RR interval (s)** | | | |
| Intercept | 16575.0 | 1 | $< 2.2 \times 10^{-16}$ |
| Acute temperature (°C) | 9514.5 | 1 | $< 2.2 \times 10^{-16}$ |
| Season | 260.4 | 1 | $< 2.2 \times 10^{-16}$ |
| Drug treatment | 55.0 | 1 | $1.23 \times 10^{-13}$ |
| Temperature x Season | 33.3 | 1 | $8.05 \times 10^{-9}$ |
| Temperature x Drug | 12.2 | 1 | $4.81 \times 10^{-4}$ |
| Season x Drug | 41.9 | 1 | $9.49 \times 10^{-11}$ |
| Temperature x Season x Drug | 16.4 | 1 | $5.14 \times 10^{-5}$ |
| **QT interval (s)** | | | |
| Intercept | 0.1 | 1 | 0.73 |
| Acute temperature (°C) | 621.2 | 1 | $< 2.2 \times 10^{-16}$ |
| Season | 0.4 | 1 | 0.51 |
| Drug treatment | 0.2 | 1 | 0.66 |
| Temperature x Season | 5.7 | 1 | 0.02 |
| Temperature x Drug | 8.3 | 1 | $4.06 \times 10^{-3}$ |
| Season x Drug | 0.4 | 1 | 0.54 |

Model: glmer(RR~Temperature*Season*Drug + (1|ID), ARCH, family=Gamma(link='log'))
Model: glmer(QT~Temperature*Season+Temperature*Drug+Season*Drug + (1|ID), ARCH, family=Gamma(link='log'))

**Table S5.** Results of the generalized linear mixed effect models for the change in diastole and diastole/systole duration under acute warming for maximum and intrinsic heart rate treatments and seasonal treatments (winter and summer) of wild juvenile Arctic char. Diastole and diastole/systole duration were modeled from 5°C to peak$f_H$. Fish ID was assigned as the random effect. Significant effects ($P < 0.05$) calculated using a type III Wald chi-square test.

| Fixed effect | Chi-square | df | *P* |
|---|---|---|---|
| **Diastole duration** | | | |
| Intercept | $2 \times 10^{-4}$ | 1 | 0.99 |
| Acute temperature (°C) | 356.9 | 1 | $< 2.2 \times 10^{-16}$ |
| Drug treatment | 2.6 | 1 | 0.11 |
| Season | 0.5 | 1 | 0.50 |
| Acute temperature x Drug treatment | 7.3 | 1 | $6.87 \times 10^{-3}$ |
| **Diastole/systole duration** | | | |
| Intercept | 4.0 | 1 | 0.04 |
| Acute temperature (°C) | 1.0 | 1 | 0.33 |
| Season | 2.8 | 1 | 0.10 |
| Drug treatment | 0.7 | 1 | 0.42 |
| Body mass (g) | 4.0 | 1 | 0.046 |
| Acute temperature x Season | 0.1 | 1 | 0.74 |
| Acute temperature x Drug treatment | 0.2 | 1 | 0.63 |
| Season x Drug | 5.8 | 1 | 0.02 |
| Acute temperature x Season x Drug | 6.9 | 1 | $8.53 \times 10^{-3}$ |

Model: glmer(Diastole ~Temperature*Drug + Season + (1|ID), ARCHFull, family = Gamma(link = "log"))
Model: glmer(DS ~ Temperature * Season * Drug + Mass + (1|ID), ARCHFull, family=Gamma("log"))

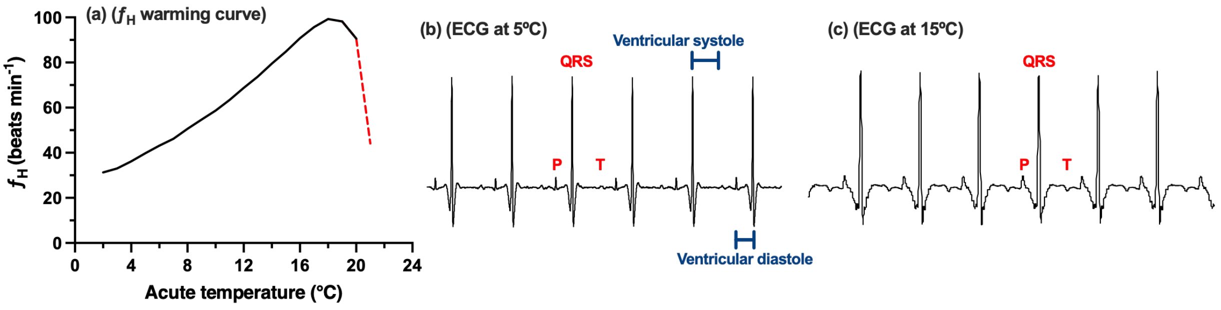


**Figure S1**. The response of heart rate ($f_H$) to acute warming and changes seen in the electrocardiogram. (a) The $f_H$ over a period of warming with $f_H$ after cardiac arrhythmia shown in red. (b) Electrocardiogram tracings of a winter fish at 5°C and (c) a summer fish at 15°C. Waveform components are identified on the ECG (P; atrial depolarization, QRS; ventricle depolarization; T, ventricle repolarization).

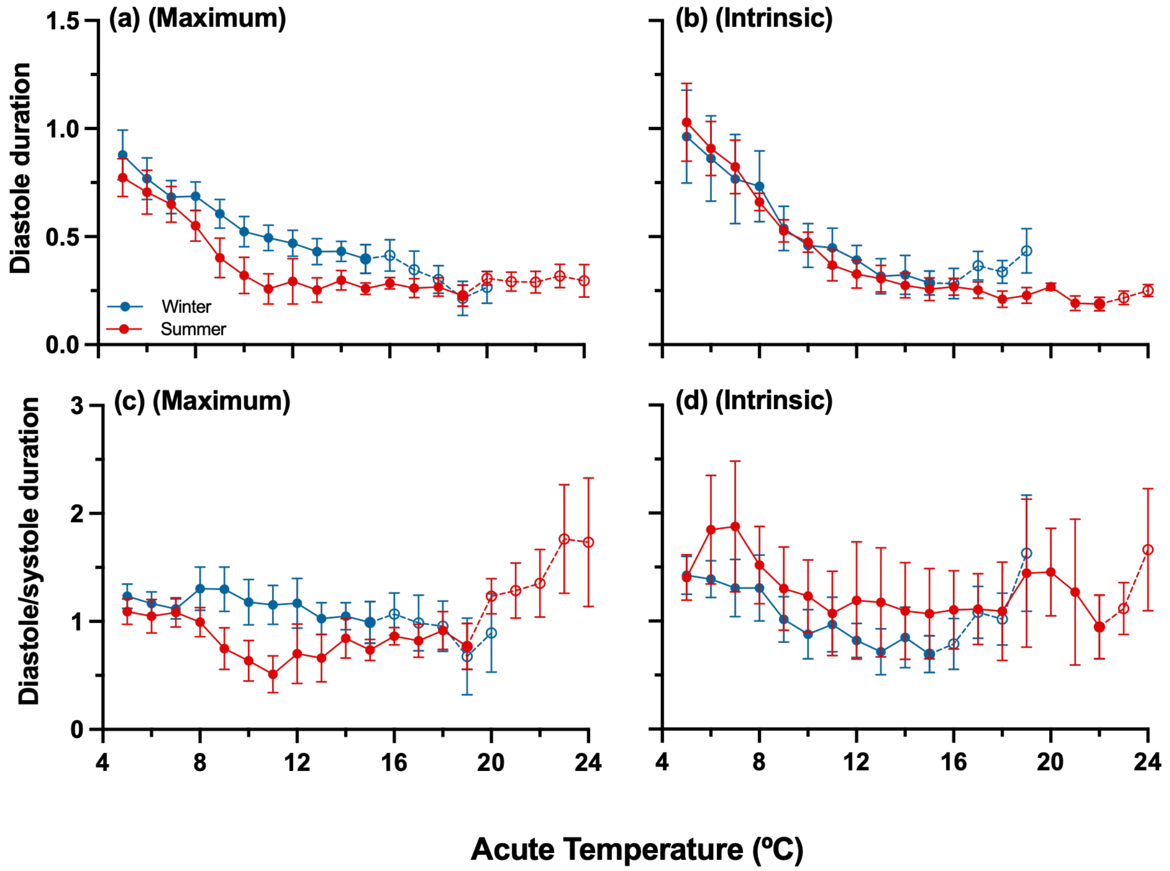


**Figure S2.** Effect of acute warming on seasonally acclimatized juvenile Arctic char contraction and relaxation periods of the cardiac cycle. Diastole duration (a, b) of seasonally acclimatized fish for maximum (a) and intrinsic (b) treated fish. Diastole/systole (c, d) of seasonally acclimatized fish for maximum (c) and intrinsic (d) treated fish. Estimated acclimatization temperatures (average body temperature during the experimental periods) were 1.1 ºC in winter (n=21) and 11.1 ºC in summer (n=18).

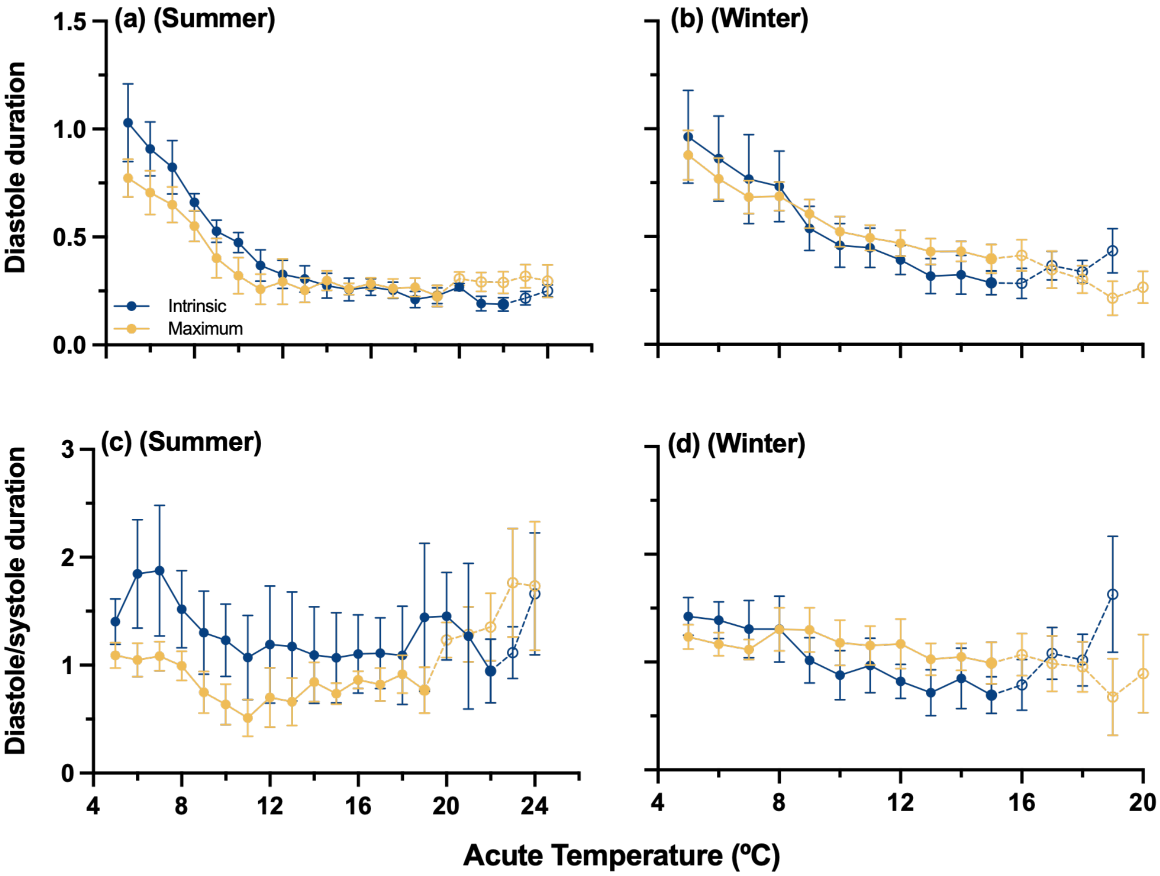


**Figure S3.** Effect of acute warming on seasonally acclimatized juvenile Arctic char contraction and relaxation periods of the cardiac cycle. Diastole duration (a, b) of maximum and intrinsic treated fish for summer (a) and winter (b) acclimatized fish. Diastole/systole (c, d) of maximum and intrinsic treated fish for summer (c) and winter (d) acclimatized fish. Acclimatization temperatures (average body temperature during the experimental periods) were 1.1 ºC in winter (n=21) and 11.1 ºC in summer (n=18).

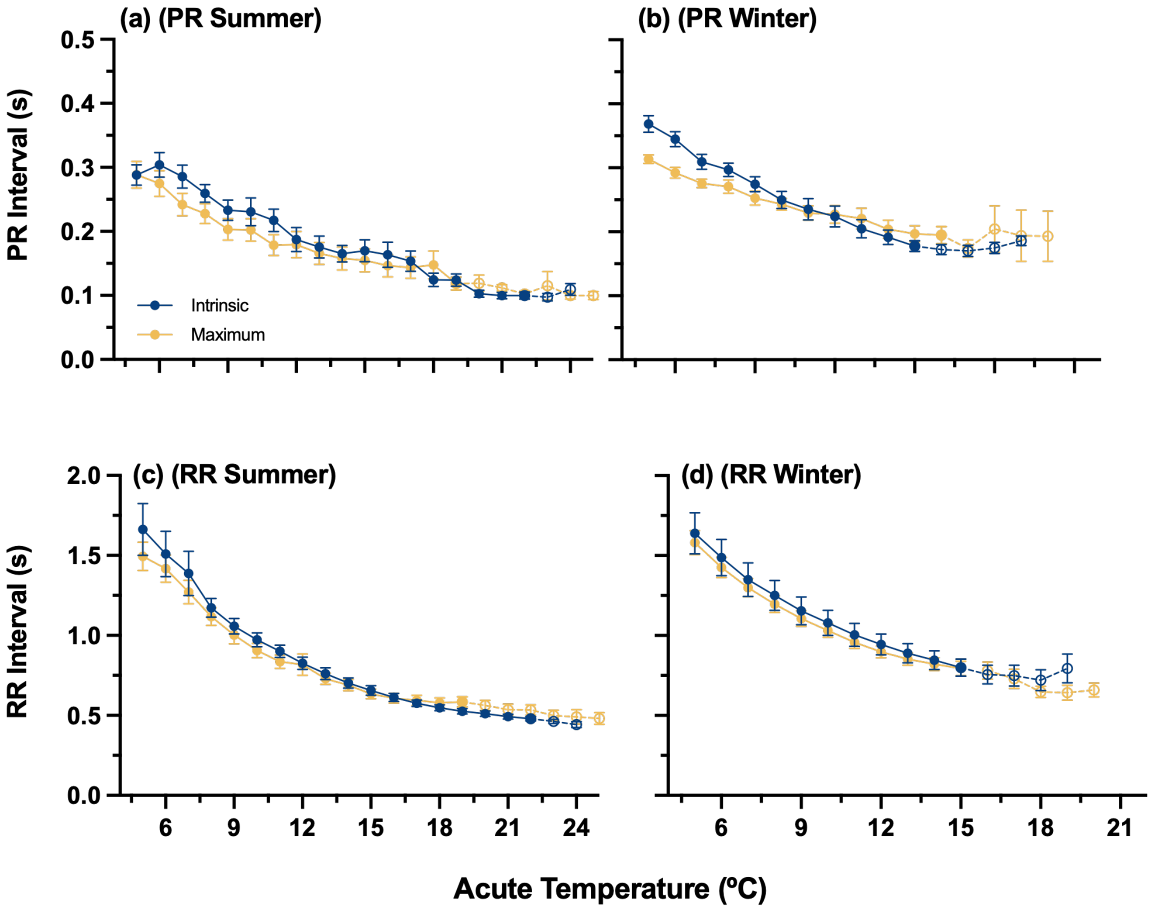


**Figure S4.** Effect of acute warming and drug treatment on the PR interval (a, b), and RR interval (c, d) for summer acclimatized (a, c) and winter acclimatized (b, d) fish's heart rates. Data are presented as mean ± SEM, open circles and dashed lines indicate where fish were removed after onset of arrhythmia. Acclimatization temperatures (average body temperature during the experimental periods) were 1.1 ºC in winter (n=21) and 11.1 ºC in summer (n=18).

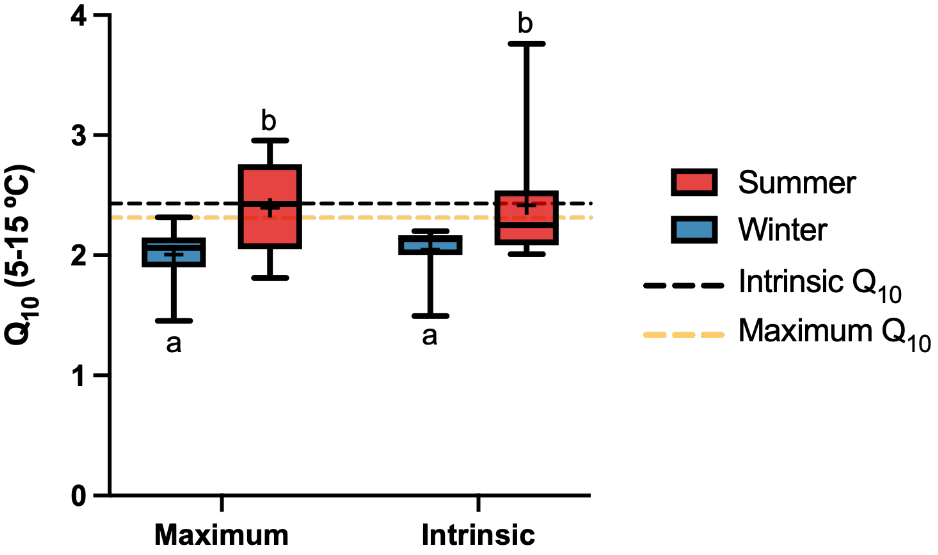


**Figure S5**. Effect of seasonal acclimatization and drug treatment on the thermal sensitivity ($Q_{10}$) of heart rate between 5 and 15 ºC in seasonally acclimatized juvenile Arctic char. Box plots represent $Q_{10}$ values calculated from individual acute values where a $Q_{10}$ value was calculated for individuals between 5 and 15 ºC. Dashed lines show the acclimated $Q_{10}$ values calculated by averaging all winter acclimatized $f_H$ values at 5 ºC and all summer acclimatized $f$H values at 15 ºC and calculating a $Q_{10}$ value between those two averages. Acute and acclimated $Q_{10}$ values were calculated by drug treatment. Boxes represent median and interquartile ranges, with whiskers as the 5$^{th}$ and 95$^{th}$ data percentile, crosses indicate the mean.
